# Comprehensive Characterization of Toxicity of Fermentative Metabolites on Microbial Growth

**DOI:** 10.1101/169482

**Authors:** Brandon Wilbanks, Cong T. Trinh

**Affiliations:** Department of Chemical and Biomolecular Engineering, University of Tennessee, Knoxville, TN 37920; Bioenergy Science Center (BESC), Oak Ridge National Laboratory, Oak Ridge

**Keywords:** *Escherichia coli*, toxicity, alcohols, carboxylic acids, esters, growth inhibition, energy density, partition coefficient, hydrophobicity

## Abstract

**Background:** Volatile carboxylic acids, alcohols, and esters are natural fermentative products, typically derived from anaerobic digestion. These metabolites have important functional roles to regulate cellular metabolisms and broad use as food supplements, flavors and fragrances, solvents, and fuels. Comprehensive characterization of toxic effects of these metabolites on microbial growth under similar conditions is very limited.

**Results:** We characterized a comprehensive list of 32 short-chain carboxylic acids, alcohols, and esters on microbial growth of *Escherichia coli* MG1655 under anaerobic conditions. We analyzed toxic effects of these metabolites on *E. coli* health, quantified by growth rate and cell mass, as a function of metabolite types, concentrations, and physiochemical properties including carbon chain lengths and associated functional groups, chain branching features, hydrophobicity, and energy density. Strain characterization reveals these metabolites exerted distinct toxic effects on *E. coli* health. We find that higher concentrations and/or longer carbon lengths of metabolites cause more severe growth inhibition. For the same carbon lengths and metabolite concentrations, alcohols are most toxic followed by acids then esters. We also discover that branched chain metabolites are less toxic than linear chain metabolites for the same carbon lengths and metabolite concentrations. Remarkably, shorter alkyl esters (e.g., ethyl butyrate) are found to be less toxic than longer alkyl esters (e.g., butyl acetate) for the same carbon lengths and metabolite concentrations. Regardless of metabolite types, longer chain metabolites are less soluble and have higher energy densities but are more toxic to microbial growth.

**Conclusions:** Metabolite hydrophobicity, correlated with carbon chain length, associated functional group, chain branching feature, and energy density, is a good quantitative index to evaluate toxic effect of a metabolite on microbial health. The results provide better understanding of degrees of toxicity of fermentative metabolites on microbial growth and further help selection of desirable metabolites and hosts for industrial fermentation to overproduce them.

## BACKGROUND

During anaerobic digestion of organic matters, organisms naturally produce volatile organic acids and alcohols to balance cellular redox states. These molecules, along with esters generated from condensation of alcohols and acids, are of particular interest for not only fundamentally studying their functional roles to regulate cellular metabolisms and microbiomes [1] but also harnessing them as food supplements, natural flavors and fragrances, solvents, and fuels [2].

A diverse class of microbes can naturally produce these volatile metabolites, and some being harnessed for industrial-scale production. For instance, *Escherichia coli*, a facultative, gram-negative bacterium found in lower intestine of animals, is widely used as an industrial workhorse microorganism for biocatalysis. *E. coli* possesses a native mixed acid fermentative metabolism that has been metabolically engineered to produce many of fermentative metabolites including alcohols (e.g., ethanol [3, 4], isopropanol [5], butanol [6], isobutanol [7], pentanol [8], and hexanol [9]), diols (e.g., 1,3-propanediol [10], and 1,4-butanediol [11]), acids (e.g., pyruvate [12], lactate [13], and short-medium chain carboxylic acids [14]), diacids (e.g., succinate [15], adipate [16]), and esters (e.g., acetate esters [17], propionate esters [18, 19], butyrate esters [18-20], pentanoate esters [18, 19], and hexanoate esters [18, 19]).

Fermentative metabolites, however, can become inhibitory to microbial growth by directly interfering with cell membrane and/or intracellular processes [21-29]. Currently, data on toxic effects of a comprehensive set of fermentative metabolites on microbial growth under similar growth conditions is very limited. Availability of this data can help identify and better understand most toxic metabolites to microbes during fermentation. It also provides design criteria for selecting desirable metabolites and microbes for industrial production as well as effective engineering strategies to alleviate toxicity. For instance, various strategies of targeted engineering have been implemented to enhance microbial tolerance against some fermentative metabolites including increasing the ratio of saturated and unsaturated fatty acid compositions [30], raising the average chain length of fatty acid moieties in cell membrane [31], enhancing the ratio of trans- and cis-unsaturated fatty acids of cell membrane [32], and expressing efflux pumps [33] or chaperones [34]. Genome and evolutionary engineering have also been explored to enhance tolerance [24, 35-37].

In this study, we characterized toxic effects of a comprehensive set of 32 fermentative metabolites including 8 carboxylic acids, 8 alcohols, and 16 esters on *E. coli* health. We analyzed toxic effects of these metabolites as a function of metabolite types, concentrations, and physiochemical properties including carbon chain lengths and associated functional groups, chain branching features, hydrophobicity, and energy density.

## RESULTS AND DISCUSSION

To study toxic effects of fermentative metabolites on *E. coli* health, growth kinetics were generated for each metabolite using standard concentrations (0, 2.5, 5.0, and 7.5 g/L) and additional concentrations as needed for certain metabolites. Both maximum growth rate and optical density (OD) during the first 24 h period were extracted to evaluate *E. coli* health. For the reference growth condition without supplementation of a toxic chemical, wildtype *E. coli* MG1655 grew at a rate of 0.6134 ± 0.0337 1/h and OD of 1.3982 ± 0.0554 (Figure 1).

**Figure 1:**
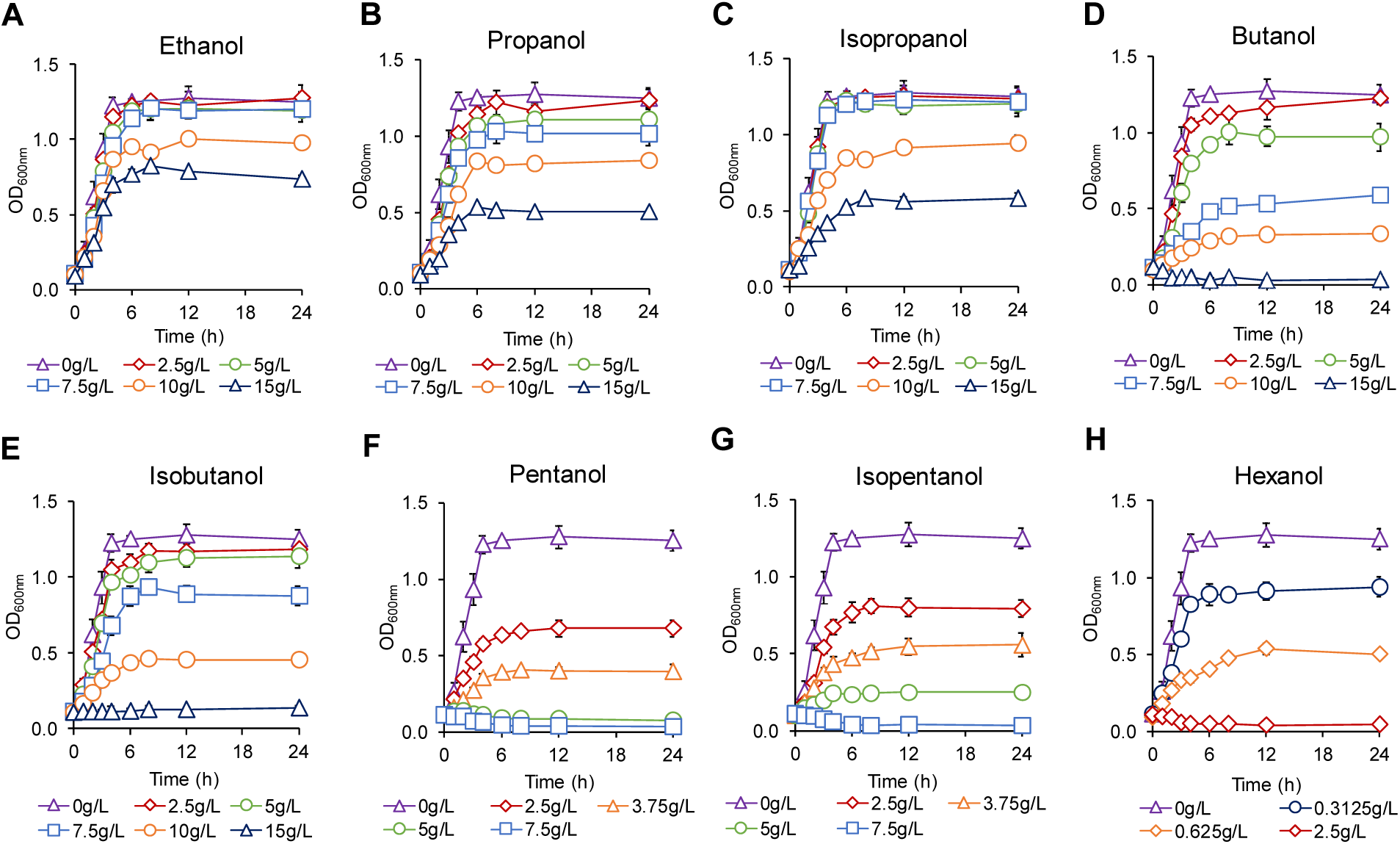
Growth kinetics of *E. coli* exposed to eight short-chain alcohols at various concentrations including **(A)** ethanol, **(B)** propanol, **(C)** isopropanol, **(D)** butanol, **(E)** isobutanol, **(F)** pentanol, **(G)** isopentanol, and **(H)** hexanol.

### Toxic effects of alcohols

The first alcohol of interest, ethanol, was found to be essentially non-toxic up to 7.5 g/L (Figure 1A). At 10 g/L ethanol, specific growth rate and OD decreased by only 12% and 25% each as compared to the reference (without supplementation of the toxin) (Figure 2). At the highest measured concentration of 15 g/L, growth rate was further reduced by only 18%, but OD was nearly 40% lower at 0.8240 ± 0.0130.

**Figure 2:**
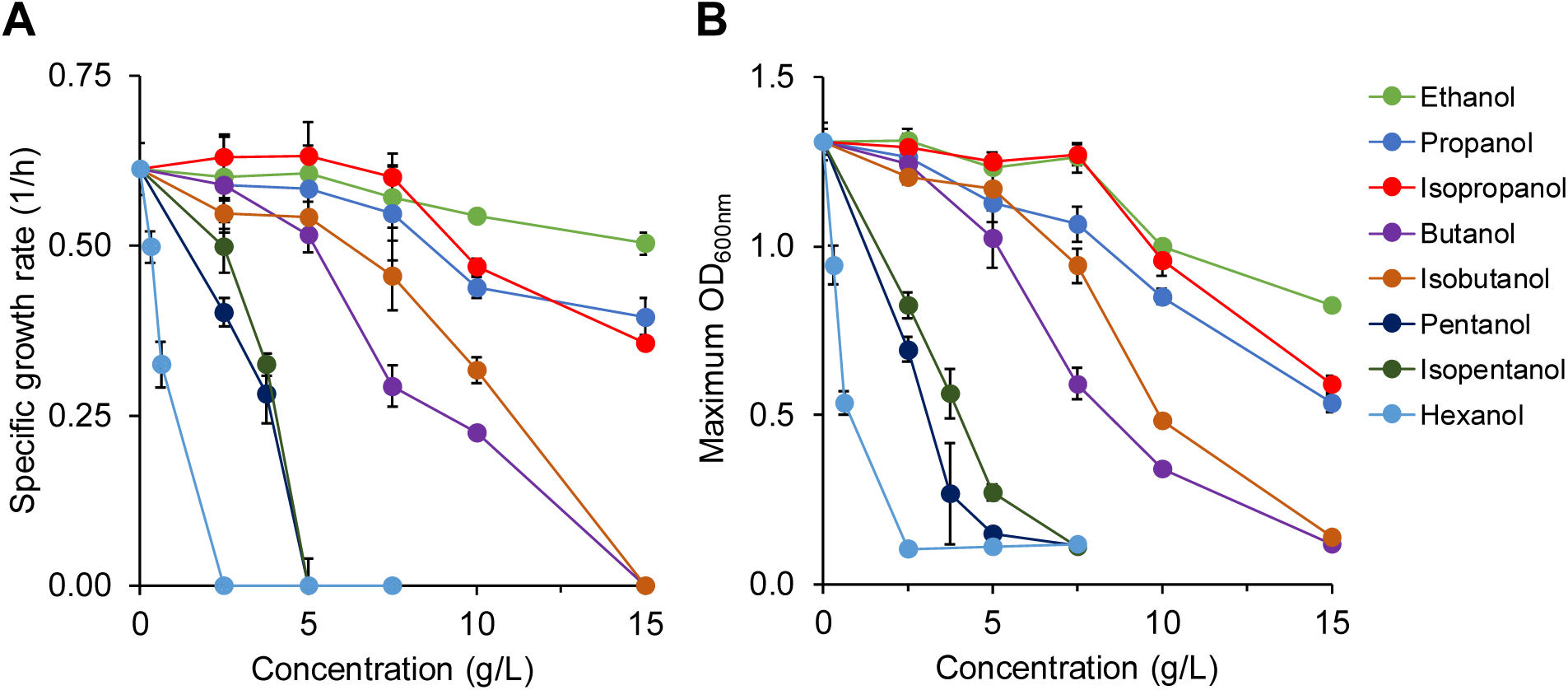
Comprehensive analysis of toxic effects of alcohols on *E. coli* health based on **(A)** specific growth rate and **(B)** OD.

Propanol toxicity at concentrations up to 7.5 g/L was similar to that of ethanol, but at 15 g/L it was significantly more toxic (Figure 1B). Specific growth rate was 0.3955 ± 0.0278 1/h (nearly 50% lower than the reference) and OD was 0.5337 ± 0.0271 (∼60% lower than the reference) (Figure 2). Isopropanol toxicity exhibited relatively similar trends like propanol toxicity but with slightly higher growth rate and OD at most concentrations tested (Figures 1C, 2).

Butanol is the first alcohol to display strong toxic effects before 10 g/L (Figure 1D). At 7.5 g/L, growth rate (0.2932 ± 0.0302 1/h) and OD (0.5927 ± 0.0454) were reduced more than 50% as compared to the reference (Figure 2). Growth was entirely inhibited in butanol at 15 g/L. Our data presented for butanol toxicity is consistent with a previous study reporting that growth of *E. coli* DH5α in YPD medium was reduced by 80% in 1% v/v (∼8.1 g/L) butanol and stopped at 2% v/v (∼16.2 g/L) [38]. Isobutanol was less toxic than butanol at all concentrations, with the exception of 15 g/L, where no growth was observed for both compounds (Figure 1E). At 7.5 g/L, isobutanol was less inhibitory than butanol for *E. coli* growth, with higher specific growth rate and OD by approximately 25% (Figure 2). Findings of isobutanol toxicity presented here are consistent with the Atsumi *et al*.’s report [24]. The difference in toxic effects of isobutanol and butanol is consistent with the data by the Huffer *et al.* report [25]. Remarkably, based on the Huffer *et al*.’s data, microbial health is less inhibited in isobutanol than butanol for not only *E. coli* but also some other bacterial, eukaryotic, archaeal species.

For pentanol and isopentanol, no growth was observed at any studied concentrations above 5 g/L (Figures 1F, 1G). Pentanol terminated all growth at 5g/L, and at 3.75 g/L specific growth rate was just 0.2818 ± 0.0438 1/h (Figures 1F, 2). Unlike pentanol, isopentanol at 5 g/L allowed for growth, with a significantly reduced specific growth rate of 0.2017 ±0.0388 1/h and a OD of 0.2703 ± 0.0241 (Figures 1G, 2). At 2.5 g/L, isopentanol suppressed specific growth rate and OD by 12% and 8% less than did pentanol.

Hexanol is the most toxic among alcohols used in this study. It eliminated all growth at only 2.5 g/L. A far reduced concentration of 0.625 g/L still cut specific growth rate by over 45% and OD by nearly 60% as compared to the reference (Figures 1H, 2).

Overall, alcohols are toxic to microbial growth, and degrees of toxicity depend on alcohol types and concentrations. Increasing alcohol concentrations decrease both specific growth rate and OD. Shorter chain length alcohols (ethanol, propanol, isopropanol) require higher concentrations in order to impact growth significantly.

### Toxic effects of carboxylic acids

Acetic acid was marginally toxic up to 7.5 g/L, at which specific growth rate (0.4392 ± 0.0320 1/h) and OD (0.9050 ± 0.0131) were each reduced by ∼20% compared to the reference (Figures 3A, 4). Propionic acid at an identical concentration was found to be much more toxic than acetic acid, with specific growth rate (0.2374 ± 0.0253 1/h) and OD (0.3542 ± 0.0142) decreased ∼60% and ∼75%, respectively (Figures 3B, 4).

**Figure 3:**
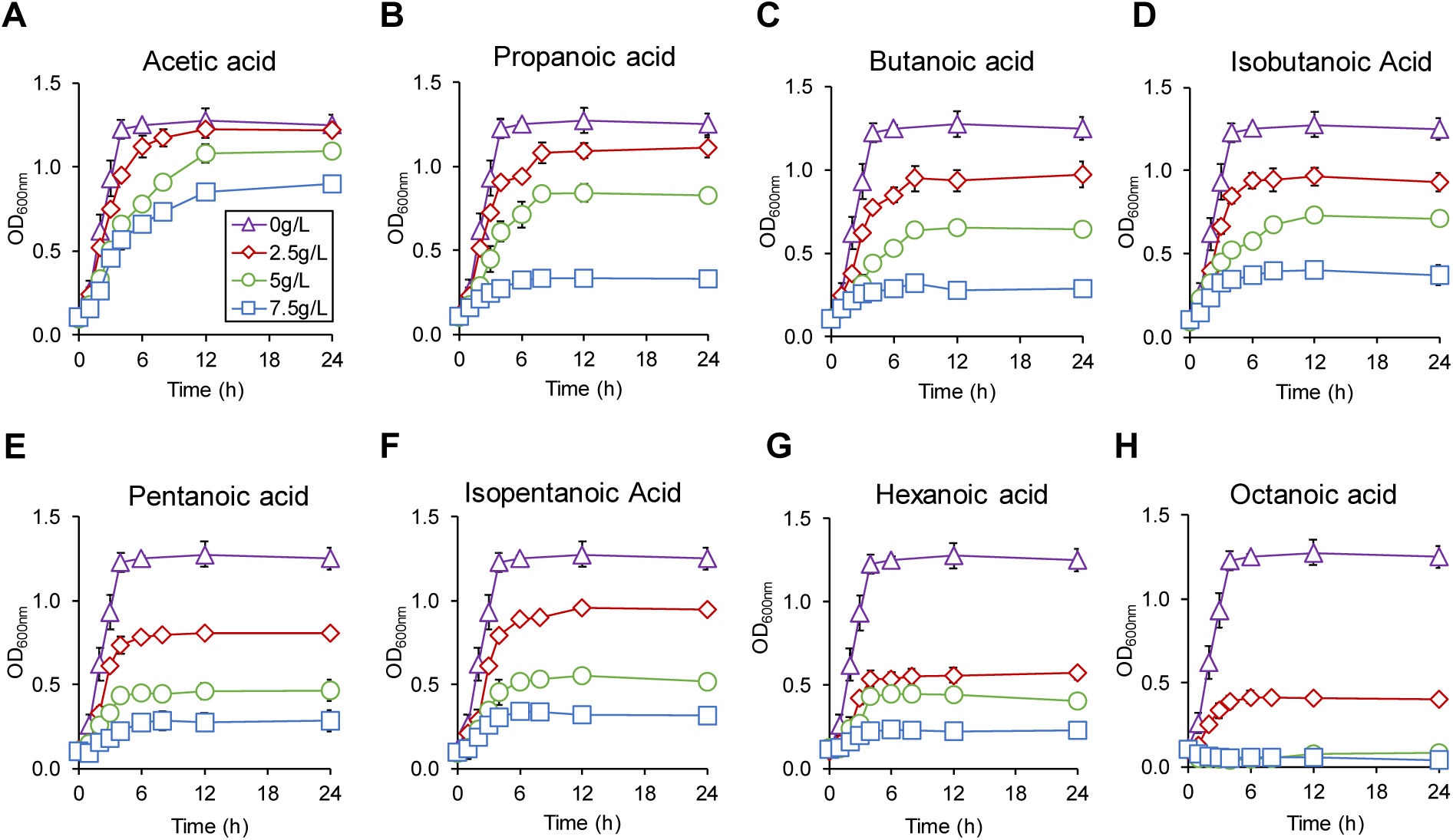
Growth kinetics of *E. coli* exposed to eight short-chain fatty acids at various concentrations including **(A)** acetic acid, **(B)** propanoic acid, **(C)** butanoic acid, **(D)** isobutanoic acid, **(E)** pentanoic acid, **(F)** isopentanoic acid, **(G)** hexanoic aid, and **(H)** octanoic acid.

**Figure 4:**
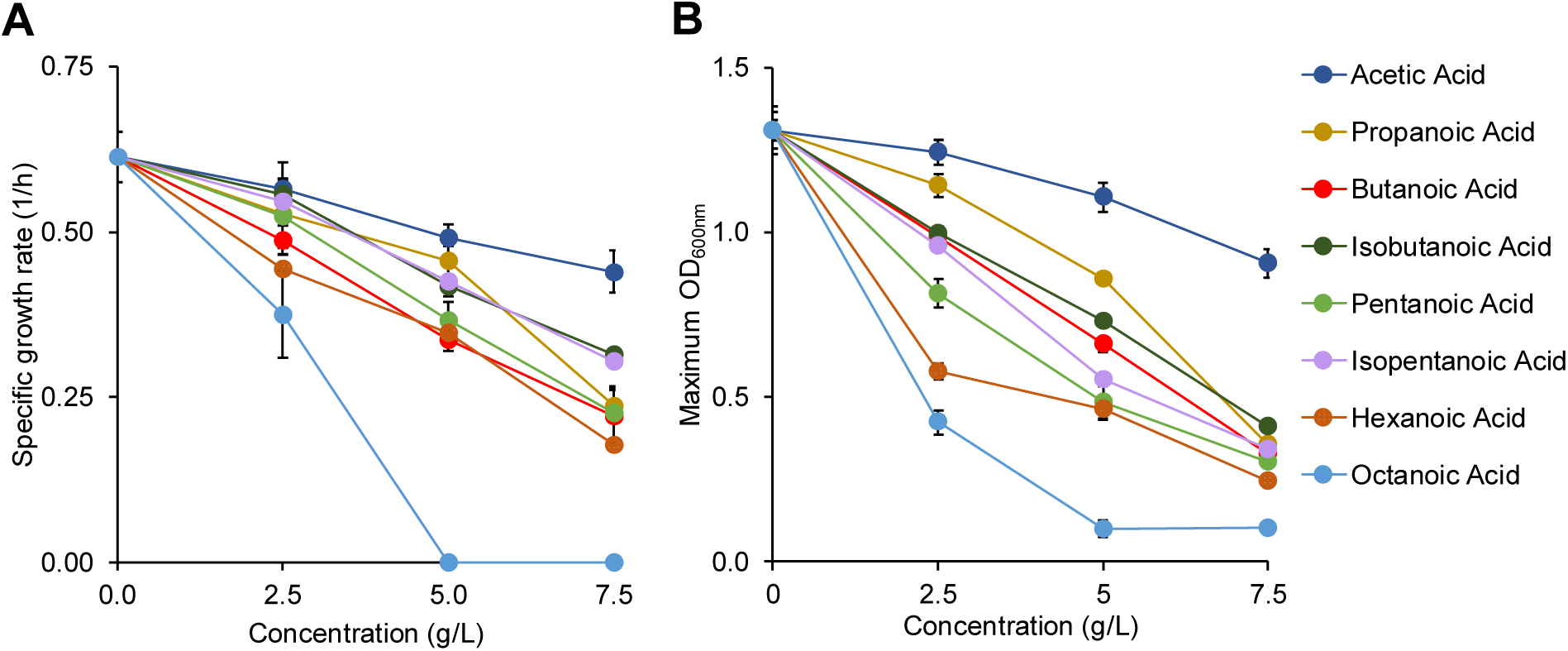
Comprehensive analysis of toxic effects of alcohols on *E. coli* health based on **(A)** specific growth rate and **(B)** OD.

Butanoic acid at 7.5 g/L was seen to be slightly more inhibitive of specific growth rate and OD than propionic acid whereas concentrations of 2.5 g/L and 5 g/L appeared similarly toxic like propionic acid (Figures 3C, 4). Isobutanoic acid was found to be less toxic than butanoic acid, following the chain branching trend seen in alcohols (Figures 3D, 4). At 2.5, 5.0, and 7.5 g/L, cells grew 6%, 5%, and 15% faster in isobutanoic acid than butanoic acid.

The pair of pentanoic and isopentanoic acid was also used. At each concentration, isopentanoic was less toxic than pentanoic acid. Pentanoic and isopentanoic acids sustained growth at 7.5 g/L to ODs of 0.3017 ± 0.0504 and 0.3417 ±0.0213, respectively, and specific growth rates reached 0.2262 ± 0.0395 and 0.3041 ± 0.0170 1/h, respectively (Figures 3E, 3F, 4).

The next acid studied was hexanoic acid. Growth with this compound was sustained at 7.5 g/L, but specific growth rate was reduced by □70% and OD just reached 0.2448 ± 0.0283 (Figures 3G, 4). Octanoic acid was even more toxic, eliminating all growth at 5 g/L (Figure 3H, 4). At 2.5 g/L, specific growth rate (0.3741 ± 0.0598 1/h) and OD (0.4328 ± 0.0219) was decreased by about 40% and 65% as compared to the reference, respectively. Octanoic acid is the most toxic organic acid studied here, and the only acid that prevented all growth above 2.5 g/L.

Like alcohols, acid toxicity on microbial growth depends on exposed concentrations and acid types. Increasing acid concentrations enhances toxicity for all compounds, reducing growth rates and cell concentrations. Longer chain acids cause more severe growth inhibition

### Toxic effects of esters

Cells can produce a combinatorial library of esters by condensing organic acids and alcohols [18-20]. In this study, we investigated the toxic effects of a comprehensive list of 16 common short-chain esters on *E. coli* health. For comparison, we classified these esters into 3 categories: ethyl esters, propyl esters, and butyl esters.

#### Ethyl esters

Ethyl acetate was not strongly toxic until concentrations of 10 g/L or greater (Figure 5A). At 10 and 15 g/L, specific growth rates observed were reduced to 0.4246 ± 0.0089 1/h and 0.2664 ±0.0073 1/h, respectively. OD followed similar trends, being reduced to 0.8677 ± 0.0311 at 10 g/L and 0.3490 ± 0.0255 at 15 g/L (Figure 6). Ethyl propionate was more toxic than ethyl acetate at identical concentrations (Figure 5B). At 10 g/L, specific growth rates between growth in ethyl acetate and ethyl propionate were not significantly different, but OD was more than 20% lower in ethyl propionate than in ethyl acetate (Figure 6). No growth occurred with the addition of 15 g/L ethyl propionate, making ethyl acetate the only ester that allowed for any growth at 15 g/L (Figure 5).

**Figure 5:**
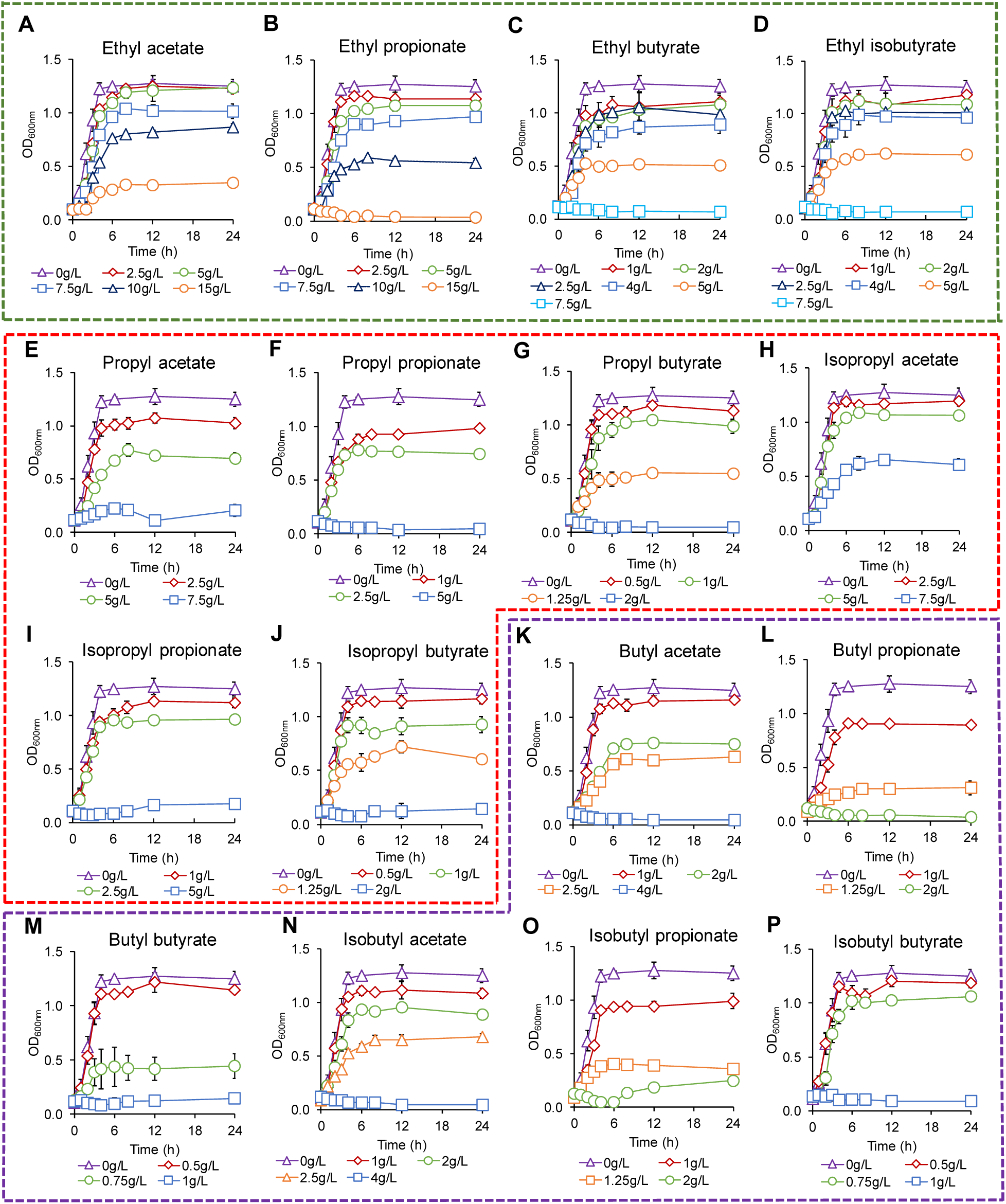
Growth kinetics of *E. coli* exposed to sixteen short-chain esters at various concentrations including **(A-D)** ethyl esters, **(E-J)** (iso)propyl esters, and **(K-P)** (iso)butyl esters

**Figure 6:**
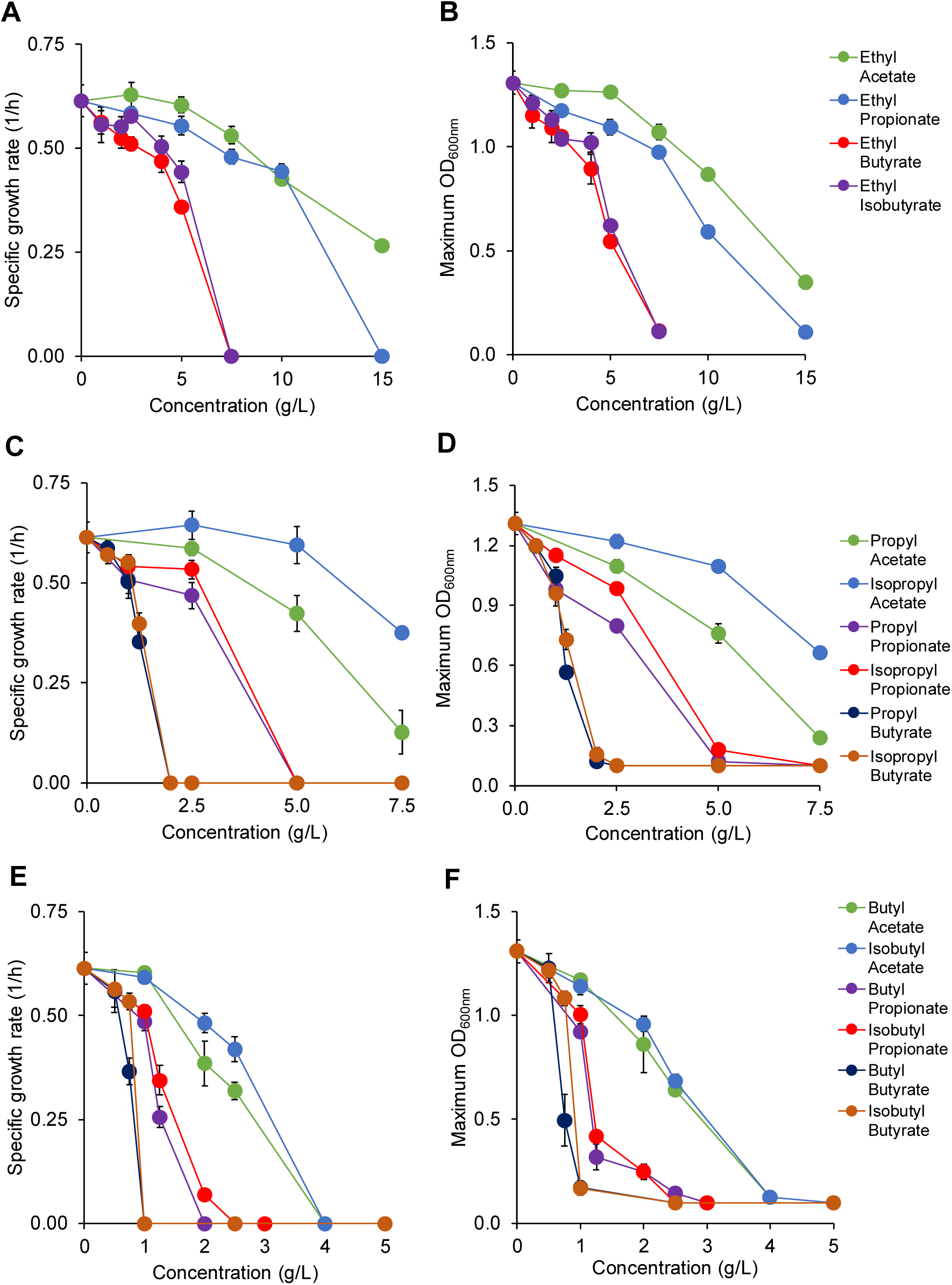
Comprehensive analysis of toxic effects of esters on *E. coli* health based on specific growth rate and OD of **(A-B)** ethyl esters, **(C-D)** (iso)propyl esters, and **(E-F)** (iso)butyl esters.

Ethyl butyrate was the most toxic among the characterized ethyl esters, with a specific growth rate of 0.3592 ± 0.0050 1/h and OD of 0.5437 ± 0.0151 at 5 g/L (Figures 5C, 6). The toxic effect of ethyl butyrate was still noteworthy at 5 g/L, slowing growth rate by over 25% and lowering OD by over 40% as compared to the reference. The branched chain isomer of ethyl butyrate, ethyl isobutyrate, was also studied (Figure 5D). It was less toxic than ethyl butyrate at all concentrations, most notably at 5 g/L, where observed growth rate was approximately 20% higher than the growth rate with ethyl butyrate (Figure 6). Cultures with 7.5 g/L of both ethyl butyrate and ethyl isobutyrate were unable to grow (Figures 5C, 5D).

#### Propyl esters

Both propyl acetate and isopropyl acetate inhibited growth at 7.5 g/L, but isopropyl acetate was far less toxic (Figures 5E, 5H). Cultures containing propyl acetate at 7.5 g/L reached an OD of 0.2372 ± 0.0241, doubling only once in 24 h of characterization. However, the cell culture with isopropyl acetate at 7.5 g/L displayed a higher OD than the cell culture with propyl acetate by 3 folds (Figure 6). Cells (0.3749 ± 0.0148 1/h) also grew 3.5 times faster in isopropyl acetate than propyl acetate at this concentration.

The addition of propyl propionate at any concentration 5 g/L or higher prevented all growth (Figure 5F). A strong toxic effect was seen at the addition of 2.5 g/L of the compound, reducing both specific growth rate (0.4689 ± 0.0234 1/h) and OD (0.7962 ± 0.0168) by ∼25% and ∼40% as compared to the reference, respectively (Figure 6). On the other hand, cultures exposed to 2.5 g/L isopropyl propionate displayed much healthier growth (Figures 5I, 6), with a specific growth rate of 0.5332 ± 0.0329 (1/h) and OD of 0.9837 ± 0.0209. Like propyl propionate, no growth occurred in cultures at 5 g/L isopropyl propionate.

The final pair of propyl esters characterized here is propyl butyrate and isopropyl butyrate. Both compounds prevented any growth from occurring at 2 g/L, but growth was sustained at concentrations of 1.25 g/L or lower (Figures 5G, 5J). Propyl butyrate at 1.25 g/L decreased specific growth rate (0.3527 ± 0.0077 1/h) and OD (0.5670 ± 0.0277) about 2 folds. Isopropyl butyrate was less toxic, with 7% higher growth rate and 15% higher OD than propyl butyrate at this concentration (Figure 6).

#### Butyl esters

The addition of butyl acetate reduced both specific growth rate and OD by half at a concentration of 2.5 g/L (Figures 5K, 6) while all previously discussed acetate esters (ethyl acetate, propyl acetate, isopropyl acetate) showed no toxic effects at 2.5 g/L or less. No growth was observed at any concentrations of butyl acetate higher than 4 g/L. Isobutyl acetate was less toxic than butyl acetate where cells (0.4194 ± 0.0294 1/h) grew 15% faster at 2.5 g/L and displayed a 3% increase in OD (0.6847 ± 0.0341 1/h) (Figures 5N, 6). Like butyl acetate, cells exposed to isobutyl acetate at concentrations higher than 4 g/L failed to grow.

Butyl propionate is far more toxic than butyl acetate (Figures 5L, 6). Unlike butyl and isobutyl acetates, butyl propionate with concentration greater than 2 g/L prevented growth. Growth at 1.25 g/L of this compound was marginal with specific growth rate decreased by more than 60%. The toxic effects were even seen at just 1 g/L, where specific growth rate (0.4850 ± 0.0207) dropped by 20%. Isobutyl propionate was slightly less toxic, allowing for growth at 2 g/L, but specific growth rate and OD were each no more than 20% of that of the reference (Figures 5O, 6).

The final esters of interest were the pair of butyl butyrate and isobutyl butyrate. Butyl butyrate was the most toxic compound in this work, prohibiting all growth at any concentrations of 1 g/L or higher (Figures 5M, 5P, 6). At just 0.75 g/L, specific growth rate was reduced to 0.3661 ± 0.0319 1/h (60% of the reference) and OD to 0.4948 ± 0.1426 (∼35% of the reference). In comparison, isobutyl butyrate limited growth by 30% less (Figure 6), displaying a specific growth rate of 0.5337 ± 0.0204 (1/h) at the same concentration. OD was over 2-fold higher with this compound than with butyl butyrate. Growth at concentrations of 1 g/L of both compounds was prevented.

Like alcohols and acids, we observed similar trend of toxicity as a function of ester types and concentrations. Increasing ester concentrations increases toxicity for all compounds and shorter chain esters exhibit less toxic effects on microbial growth.

There is a strong linear correlation (R^2^>0.94) between growth rate and cell mass when *E. coli* is exposed to alcohols, acids, and esters (Supplementary Figure 1). Therefore, *E. coli* health can be evaluated based on growth rates and cell mass under all conditions investigated.

### Linking physiochemical properties of metabolites and toxic effects

#### Chain length and associated functional groups

To compare toxic effects of metabolites within and across chemical classes, we first used the carbon chain length as a basis. Regardless of metabolite types and concentrations, carbon chain length was strongly correlated with growth inhibition, reducing both growth rate and cell mass (Figure 7). The longer the carbon length is, the more toxic a metabolite becomes.

**Figure 7:**
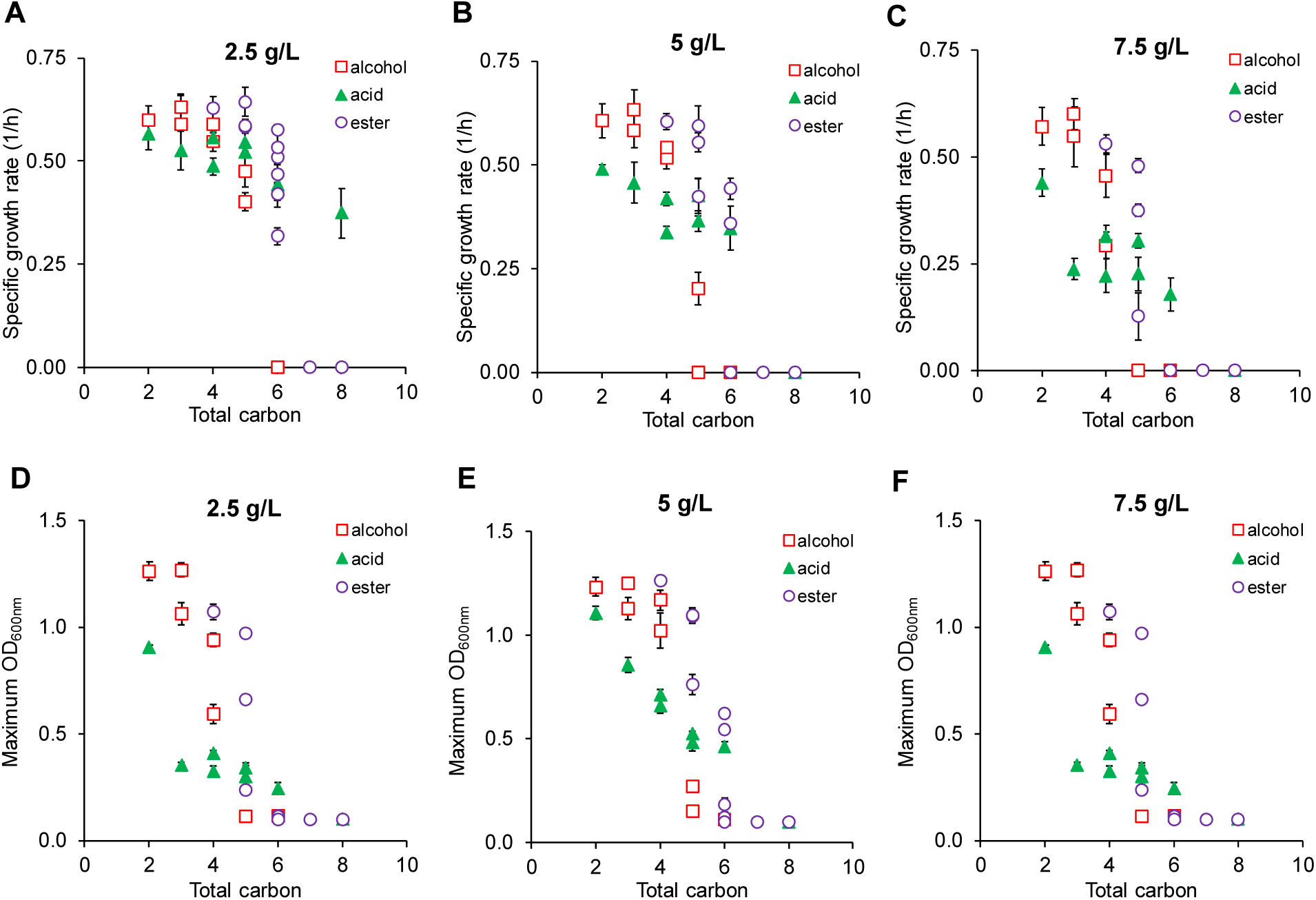
Comprehensive analysis of metabolite carbon chains determining toxic effects on *E. coli* health based on **(A-C)** specific growth rate and **(D-F)** OD at 2.5, 5.0, and 7.5 g/L metabolites.

Toxic effects of longer chain metabolites on microbial growth are likely caused by membrane disruption. Except esters, some acids and alcohols have been reported to disrupt membrane integrity and hence inhibit cell growth [25, 27, 39, 40]. As the total count of carbon atoms in a molecule increases, it becomes more soluble in the cell’s lipid membrane and less so in aqueous media. This interference causes extensive changes to cell morphology, primarily elongation due to changes in membrane fluidity, which is a well-known indicator of high stress environments and disrupted membranes [41]. This effect of chain length has been discussed in previous literature among ionic liquids [42] and surfactants [43], but has not been observed for a comprehensive set of fermentative metabolites investigated in this work.

Even though the correlation between carbon chain length and toxic effect is prevalent, the strength of this correlation varies among metabolites within and across metabolite classes. Alcohol toxicity is most strongly correlated with chain length, and each alcohol is overall more toxic than a corresponding organic acid or ester of the same total carbon. The trend, however, cannot be simply explained alone by the functional role of carbon chain length but needs to take into account of associated functional groups such as the relative polarity of a hydroxyl or carboxyl group. For example, pentanol and pentanoic acid each have the same number of carbon atoms, but pentanol is less polar (1.79 D versus 2.29 D). The higher polarity of pentanoic acid makes it less membrane-soluble than pentanol at identical concentrations, and hence is less toxic on microbial growth. Our data shows that cells grew faster in pentanoic acid (0.4016 ± 0.0212 1/h) than in pentanol (0.5228 ± 0.0519 1/h) at 2.5 g/L and yielded higher cell mass (OD = 0.8140 ± 0.0155 in pentanoic acid versus 0.6930 ± 0.0362 in pentanol). Another factor that could contribute to this difference in toxicity of alcohols and acids is stereochemistry. The larger carboxyl group on organic acids can physically hinder the acid’s ability to enter the membrane, whereas the smaller hydroxyl group will present less resistance.

#### Chain branching

For the same carbon length and chemical class, chain branching can also have different toxic effects on microbial growth. Our result shows that branched-chain isomers of each metabolite is less toxic to microbial growth across all chemical classes (Figure 7 and Supplementary Figures 2-4). This trend can be clearly seen when cells are exposed to C5 alcohols, esters, and acids. At 2.5 g/L exposure, for instance, cells grew ∼18% faster in isopentanol (0.4752 ± 0.0370 1/h) than pentanol (0.4016 ± 0.0212 1/h), 5% faster in isopentanoic acid (0.5560 ± 0.0186 1/h) than pentanoic acid (0.5528 ± 0.0519 1/h), and 10% faster in isopropyl acetate (0.6438 ± 0.0357 1/h) than propyl acetate (0.5849 ± 0.0167 1/h). For C5 acids, the trend is more significant when cells are exposed to higher concentrations. Cells grew ∼30% faster in isopentanoic acid than pentanoic acid at 7.5 g/L. The reduced toxic effects of chain branching can also be explained by the impact of membrane disruption. Branched chain isomers are less membrane soluble than their corresponding straight chain isomer at any given chain length, and hence become less toxic to microbial growth.

#### Ester dissociation

Each ester is comprised of alcohol and acid moieties. Different from alcohols and acids, toxic effects of esters can be very distinct in that different esters of the same total carbon length can have significantly different degrees of toxicity. To demonstrate, we focus on the pair of ethyl butyrate and butyl acetate (both C_5_H_12_O_2_) to examine this pattern. The difference between these two esters is that ethyl butyrate is comprised of ethanol and butyric acid moieties while butyl acetate is comprised of butanol and acetic acid moieties. At 2.5 g/L, cell grew ∼40% slower in butyl acetate (0.3186 ± 0.0207 1/h) than in ethyl butyrate (0.5106 ± 0.0168 1/h) and also yielded ∼40% lower cell mass in butyl acetate than ethyl butyrate (Figures 5, 6). This same trend is consistently seen in many other ester pairs of the same total carbon count.

This distinct toxic effect of esters can be explained by ester dissociation. Esters can be spontaneously hydrolyzed into alcohol and carboxylic acid moieties in aqueous media. At any given time, media supplied with esters for toxicity test contain some of both associated alcohols and carboxylic acids. For esters with the same carbon length, those having longer chain alcohol moieties are more toxic than those having shorter chain alcohol moieties.

#### Acid dissociation

For high carbon chain lengths and concentrations, acids appear less toxic than esters (Figure 7). For instance, at 7.5 g/L and a total carbon of 6, cells were still able to grow in acids (hexanoic acid) but neither in alcohols (hexanol) nor esters (ethyl butyrate, butyl acetate, propyl propionate, isopropyl propionate). This phenotype can be best explained by the acid dissociation that enables it to exist as the monoprotic acid and conjugate base forms. Degrees of dissociation depend on pKa of the metabolite and pH. In our experiments, the fraction of conjugate base dominated because the initial pH was adjusted to 7. Since the conjugate base is more hydrophilic than the monoprotic acid, it is less membrane soluble and hence less toxic.

#### Energy density

For biotechnological applications, energy density is one of the important physical properties. The longer the carbon chains become, the higher energy densities the metabolites contain (Supplementary Figure 5A). Among the classes of metabolites investigated in this study, alcohols have the highest energy densities followed by esters then acids with the
same chain lengths because alcohols are least oxygenated. Therefore, molecules with higher energy densities are more toxic to microbial growth.

#### Hydrophobicity

To better capture toxic effects of metabolites within and across different classes of metabolites, we further examined metabolite hydrophobicity as a basis. We used partition coefficients to determine and quantitatively compare hydrophobicity of metabolites. As expected, there is a strong linear correlation between the carbon chain lengths and partition coefficients (R^2^ ∼ 0.98) (Supplementary Figure 5B). The longer the carbon chain, the higher the partition coefficient becomes with a strong linear correlation. For the same carbon chain, chemicals may have slight differences in partition coefficients depending on associated function groups and chain branching. For instance, partition coefficients of pentanol, isopentanol, pentanoic acid, isopentanoic acid, ethyl propionate, and propyl acetate are 29.5, 15.1, 21.9, 16.2, 20.9, and 19.1, respectively.

Regardless of metabolite types and concentrations, a correlation exists between hydrophobicity of a metabolite and its toxic effect on microbial growth (Figure 8). As partition coefficients increase, negative effects on specific growth rates and ODs also increase significantly. The negative effects become severe when cells are exposed to higher chemical concentrations. Among different classes of metabolites examined in this study, alcohols are the most toxic as compared to acids and esters at the same partition coefficients and concentrations. Esters also appear to be less toxic than acids at lower partition coefficients and chemical concentrations. All compounds that prevented growth at concentrations greater than 2.5 g/L have a partition coefficient at least ∼250 times greater than ethanol. Every branched chain isomer in this work was shown to be less toxic than the associated straight chain isomer, and in each case the branched chain has a lower partition coefficient than the straight chain compound.

**Figure 8:**
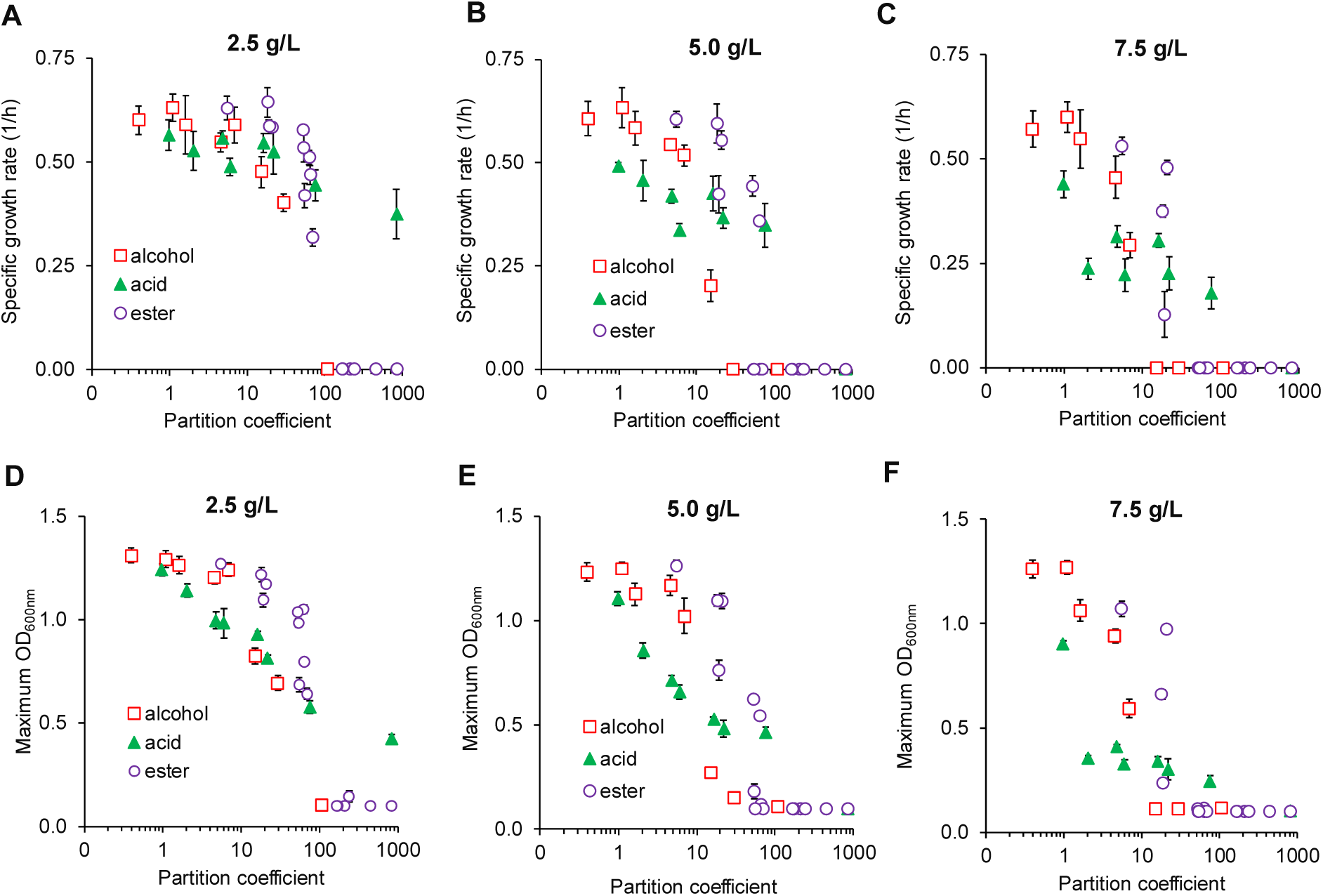
Comprehensive analysis of degrees of metabolite hydrophobicity determining toxic effects on *E. coli* health based on **(A-C)** specific growth rate and **(D-F)** OD at 2.5, 5.0, and 7.5 g/L metabolites.

Hydrophobicity of a metabolite and its toxic effect on microbial growth can be similarly explained by hydrophobic interaction between the metabolite and cell membrane. As partition coefficients increase, metabolites become more membrane soluble and disrupt lipid membranes, which enhance degrees of toxicity and sufficiently alter cell morphology [44-46]. Therefore, hydrophobicity is a good quantitative index to evaluate toxic effect of a metabolite on microbial health.

## CONCLUSION

Analysis of a comprehensive list short-chain alcohols, acids, and esters shows distinctive toxic effects of these metabolites on *E. coli* health. Alcohols are most toxic followed by acids and esters at identical concentrations and total carbon counts. Regardless of metabolite classes and concentrations, longer-chain metabolites inhibit microbial growth more significantly than shorter-chain ones. Branched-chain metabolites are less toxic than straight-chain ones with same total carbon count. Remarkably, for the same total carbon counts, esters having longer-chain alcohol moieties are more inhibitory than those having short-chain alcohol moieties. Hydrophobicity of a metabolite is a good quantitative index to determine its toxic effect on microbial health. Since this study focuses on characterizing toxic effects of fermentative metabolites on an industrial workhorse gram-negative bacterium *E. coli*, it is of particular interest to further explore in the future whether the trends found in this study exist in other bacterial, eukaryotic, and archaeal species. Even though it is not the focus of this study, fermentative metabolites can cause cytotoxicity when they are present inside the cells [23, 24, 47]. Overall, the results of this study shed light on toxic effects of fermentative metabolites with distinct characteristics on microbial growth as wells as help selection of desirable metabolites and hosts for industrial fermentation to overproduce them.

## MATERIALS AND METHODS

### Medium and cell culturing

For all *E. coli* MG1655 (DE3) characterization experiments, modified M9 medium (pH∼7) was used, consisting of 100 mL/L of 10X M9 salts, 1 ml/L of 1 M MgSO_4_, 100 μL/L of 1M CaCl_2_, 1 ml/L of stock thiamine HCl solution (1 g/L), 1 ml/L of stock trace metals solution, 10 g/L glucose, and 5 g/L yeast extract [48]. 10X M9 salts are composed 70 g/L Na_2_HPO_4_•H_2_O, 30 g/L KH_2_PO_4_, 5 g/L NaCl, and 10 g/L NH_4_Cl. Alcohols, esters, and acids were added at necessary concentrations into flasks of partitioned media. Media with the chemical of interest was then transferred from these flasks to 28 mL balch tubes and capped with rubber stoppers and aluminum seals. After addition of each chemical, media were pH adjusted to 6.5-7 with 5M KOH. Alcohols, acids, and esters were studied at varying concentrations based on a combination of factors including solubility and observed toxicity.

Stock cells from the -80^o^C freezer were struck onto lysogeny broth (LB)-agar plates and then were grown overnight in flasks containing 50 mL of the modified M9 medium in a New Brunswick Excella E25 incubator at 37^o^C and 175 rpm until OD_600nm_ (optical density measured at 600 nm using a Thermo Scientific Genysys 30 Visible Spectrophotometer) reached 2.5-3.0. In the event that this OD setpoint was surpassed, cells were diluted in 50 mL of the same medium to OD = 1.0 and grown once again to OD = 2.5. Cells were transferred to nitrogen sparged, anaerobic culture balch tubes containing 20 mL of media at initial OD = 0.1 to begin growth characterization on a 75^o^ angled platform under identical conditions. All experiments were performed in at least 3 biological replicates.

### Data collection and analysis

#### Partition coefficient

Partition coefficient, a measure of hydrophobicity of a metabolite, is calculated as follows:

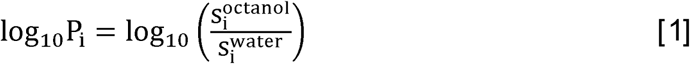

where P_i_ is the partition coefficient of metabolite i; S_i_^octanol^ and S_i_^water^ (g/L) are the solubilities of metabolite i in octanol and water, respectively. P_i_ is calculated using the Molinspiration Cheminformatics interactive log(P) calculator [49]. The input for this calculator uses the SMILES chemical notation acquired from PubChem [50].

#### ONMED

Octane Normalized Mass Energy Density (ONMED) is calculated as a ratio of standard heat of combustion of a metabolite to that of octane (∼44.5 kJ/kg) [18] where the standard heat of combustion of each chemical was estimated based on average bond energies [51].

#### Specific growth rate

First-order kinetics is applied to calculate a specific growth rate from kinetic measurement of cell growth as follows:

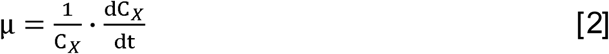

where μ (1/h) is the specific growth rate, C_X_ (g/L) is cell titer, and t (h) is culturing time. Note that in our study cell titer is estimated from measured OD with a correlation of 1 OD ∼ 0.5 g DCW/L.

## ABBREVIATIONS

μ: specific growth rate
C_X_: cell concentration
DCW: dry cell weight
OD: optical density
ONMED: octane normalized mass energy density
P_i_: partition coefficient of metabolite i
S_i_^octanol and S_i_^water^^: solubilities of metabolite i in octanol and water respectively
t: time
h: hour

## ETHICAL APPROVAL AND CONSENT TO PARTICIPATE

Not applicable.

## CONSENT FOR PUBLICATION

The authors consent for publication

## AVAILABILITY OF SUPPORTING DATA

Not applicable.

## AUTHOR’S CONTRIBUTIONS

CTT conceived and supervised the study. CTT and BW designed experiments, analyzed the data, and drafted the manuscript. BW performed the experiments. The authors have read and approved the manuscript.

## ACKNOWLEDGEMENT

We would like to thank Dr. Seunghyun Ryu for proofreading the manuscript and providing comments.

## SUPPMENTARY FILES

**Supplementary File S1**: Supplementary Figures and descriptions in a PDF.

